# The Spiking Tolman-Eichenbaum Machine: Emergent Spatial and Temporal Coding through Spiking Network Dynamics

**DOI:** 10.1101/2025.10.16.682754

**Authors:** Daisuke Kawahara, Shigeyoshi Fujisawa

**Affiliations:** RIKEN Center for Brain Science, Saitama, Japan; Department of Complexity Science and Engineering, The University of Tokyo, Tokyo, Japan

## Abstract

The hippocampal–entorhinal system supports spatial navigation and memory by orchestrating the interaction between grid cells and place cells. While various models have reproduced these patterns, many rely on predefined connectivity or fixed weights and lack mechanisms for learning or biologically realistic temporal dynamics. The Tolman–Eichenbaum Machine (TEM) has recently gained attention as a unified generative model that explains the emergence of both grid and place cells through learning. However, existing TEM implementations rely on rate-based units and simplified architectures, which limit their biological plausibility. Here, we introduce the Spiking Tolman–Eichenbaum Machine (Spiking TEM) — a spiking neural network model that extends the original TEM with spike-based computation and an anatomically inspired hippocampal–entorhinal architecture. Our model learns grid-like codes in the entorhinal module and context-specific place codes in the hippocampal module, while also exhibiting key temporal coding phenomena observed in electrophysiological recordings, including phase locking of spikes to theta oscillations and phase precession. Furthermore, the model gives rise to predictive grid cells in layer III of the entorhinal cortex, which prospectively encode upcoming spatial positions. These results demonstrate that structured spatial representations and temporally precise coding schemes can emerge from biologically plausible spike-based learning and dynamics, offering a unified framework for understanding spatial and temporal coding in the hippocampal–entorhinal circuit.

**Author Summary:** Animals, including humans, rely on an internal map of the environment to navigate and remember places. In the brain, this ability depends on two key types of neurons: place cells in the hippocampus, which fire when an animal is in a specific location, and grid cells in the entorhinal cortex, which form a hexagonal coordinate-like map of space. Understanding how neural circuits generate these spatial representations is a major goal in neuroscience.

In this study, we developed a biologically grounded spiking neural network model — the Spiking Tolman–Eichenbaum Machine (Spiking TEM)—that can learn both grid and place cell patterns through realistic neural dynamics. Our model reproduces key temporal features observed in the brain, including phase precession. In this phenomenon, neurons fire slightly earlier in each successive theta cycle as an animal moves through a place field. The model also predicts grid cells that anticipate future positions. These results provide new insight into how the hippocampal–entorhinal circuit generates and organizes spatial and memory-related representations through learning and temporal coding.

## 1 Introduction

The hippocampal–entorhinal circuit enables mammals to form structured internal representations of space, facilitating both navigation and memory. Within this circuit, grid cells in the medial entorhinal cortex (MEC) exhibit periodic spatial firing [1], while place cells in the hippocampus encode location-specific activity [2]. Understanding how these spatial codes emerge from neural dynamics remains a central challenge in neuroscience.

Several computational models have been proposed to explain the emergence of grid cells. These include oscillatory interference models [3], which rely on interference between theta-modulated inputs; continuous attractor models [4][5][6], which explain grid cells by maintaining a stable bump of activity that shifts smoothly with self-motion. The recurrent connectivity allows this activity to translate across the network, producing the periodic spatial firing patterns. While these frameworks have offered valuable insights, many rely on architectural assumptions such as hand-crafted architectures or idealized input signals—that limit their generality.

The Tolman–Eichenbaum Machine (TEM) [7] introduced a new perspective by linking spatial coding in MEC with relational memory in the hippocampus. Unlike many prior models with hand-crafted weights, TEM explains how grid-like codes can emerge as part of a general-purpose structure learning system, unifying spatial and non-spatial memory under a common computational framework. However, as a rate-based model, TEM does not incorporate temporally precise neural activity or biologically realistic mechanisms such as spike timing, synaptic dynamics, and oscillatory modulation. These temporal features are prominent in the hippocampal–entorhinal system and play a crucial role in shaping spatial codes.

In particular, phase coding—the alignment of neural spiking with specific phases of ongoing theta oscillations has been recognized as a fundamental organizing principle in hippocampal–entorhinal computation [8][9]. A hallmark manifestation of this phenomenon is phase precession, where the spikes of place cells and grid cells shift systematically to earlier phases of the theta cycle as an animal traverses their firing fields [10][11]. Phase precession provides a temporally compressed code for sequences of locations, enabling predictive representations that bridge spatial and mnemonic processing[12][13]. Despite its importance, reproducing such temporal coding features in biologically grounded models of the hippocampal–entorhinal circuit remains challenging.

To address these limitations, we propose the Spiking Tolman–Eichenbaum Machine (Spiking TEM) —a biologically grounded extension of TEM that incorporates spiking neurons, spiketiming-dependent plasticity (STDP) [14][15], and theta-modulated input [16] into its architecture. By implementing the core computational principles of TEM in a spiking neural network (SNN), we investigate whether structured spatial and temporal codes can emerge within a biologically grounded framework.

Our results demonstrate that grid-like firing patterns naturally arise in the entorhinal module of Spiking TEM alongside context-dependent place representations in the hippocampal module. Crucially, the model also exhibits phase precession in linear track simulations, showing that temporally precise coding can emerge. These codes develop through learning and are not imposed by handcrafted weights.

Furthermore, by analyzing the learned synaptic weights after training, we identify the specific structure of inputs that drive grid cell responses. This provides mechanistic insight into how spatially periodic patterns emerge through learning.

In sum, Spiking TEM bridges abstract relational modeling with spike-based neural computation, offering a unified and mechanistic account of how structured spatial and temporal codes including grid cells, place cells, and phase precession—can emerge through learning.

## 2 Methods

The source code used in the experiment is available in https://github.com/kdaisuke0203/spikingTEM.

### 2.1 Neuron model

We model each neuron using the Leaky Integrate-and-Fire (LIF) [17]. The membrane potential *V* (*t*) of a neuron evolves according to the following differential equation:

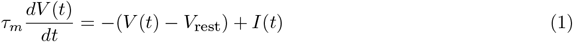

where:

- *τ*_*m*_ is the membrane time constant,
- *V*_rest_ is the resting membrane potential,
- *I*(*t*) is the total synaptic input current.

When the membrane potential reaches a predefined threshold *V*_th_, the neuron emits a spike and the membrane potential is reset:

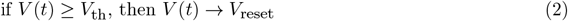

The spiking output of each neuron at discrete time *t* is represented by a binary variable *s*(*t*):

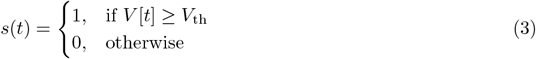

The input current *I*(*t*) to each neuron is composed of synaptic contributions from presynaptic neurons, modulated by a neuromodulatory factor *G*:

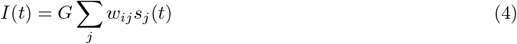

where:

- *G* is a neuromodulatory factor (e.g., representing the effects of dopamine or acetylcholine [18]) that scales the overall inputs. The neuromodulatory factors *G* were treated as learnable variables optimized through backpropagation, with initial values of 1.0. These modulations were applied to the output layers of the medial entorhinal cortex layer II (MECII) and layer III (MECIII).
- The synaptic weights *w*_*ij*_, from presynaptic neuron *j* to postsynaptic neuron *i*, were also treated as learnable variables optimized via backpropagation.
- *s*_*j*_∈(*t*) 0, 1 represents the spike at time *t* from neuron *j*.

Entorhinal and hippocampal neurons receive inhibitory inputs *I*_*θ*_ that are modulated by theta oscillations, reflecting the phase-dependent activity of local interneurons and septal GABAergic inputs [19][20][16][21]. These inputs are modeled as:

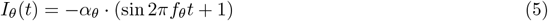

where *α*_*θ*_ *>* 0 is the amplitude and *f*_*θ*_ = 8 Hz is the theta frequency. The term (sin 2*πf*_*θ*_*t*+1) ensures that *I*_*θ*_(*t*) remains negative throughout the theta cycle, thereby maintaining a consistently inhibitory effect.

### 2.2 Network architecture

Our model consists of two interconnected components: a generative model, which predicts future sensory observations from internal latent states and actions, and an inference model, which infers latent states from current sensory observations (Fig 1), following the overall framework of the Tolman–Eichenbaum Machine (TEM) [7]. However, in contrast to the original TEM, our network architecture incorporates biologically grounded circuit motifs based on anatomical findings [22], providing a more realistic representation of the hippocampal–entorhinal system.

**Figure 1.**
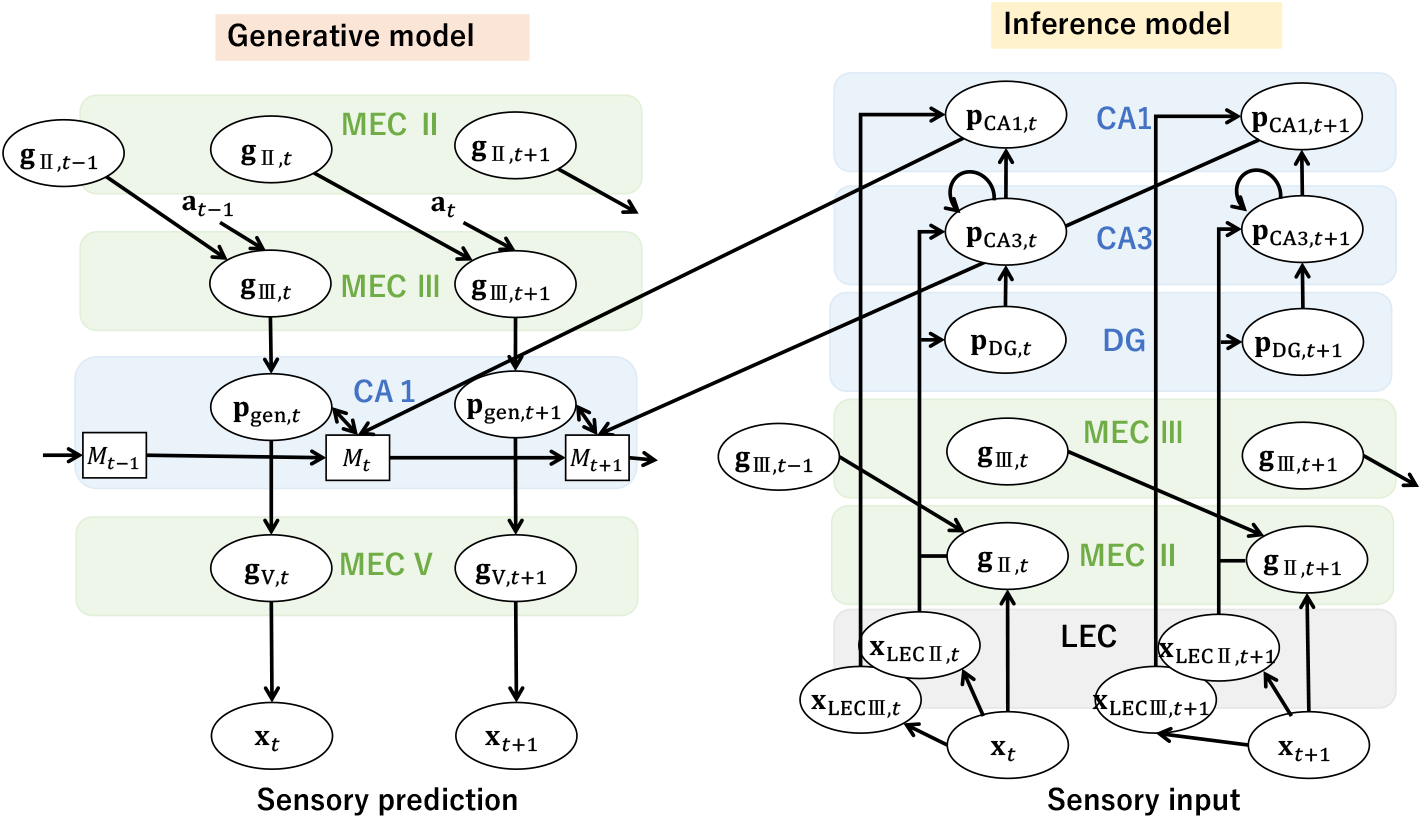
Architecture of the Spiking Tolman–Eichenbaum Machine (Spiking TEM). The model comprises two main components: a generative model (left) and an inference model (right). All computations are implemented using spiking neural networks (SNNs). EC stands for entorhinal cortex, and DG stands for dentate gyrus. *M* denotes the associative memory matrix learned via STDP. In the brain, MECII also receives input from MECV; however, in our experiments, this connection did not significantly contribute to the formation of grid cells (Table 2).

### 2.2.1 Generative model

In the generative model, the latent state **g**_ECIII,*t*_, representing neural activity in the medial entorhinal cortex (MECIII), is updated based on the previous neural activity in the MECII **g**_ECII,*t*−1_ and the previous action **a**_*t*−1_. The updated **g**_ECIII,*t*_ generates a predicted neural pattern **p**_gen,*t*_ in CA1, which, together with the memory module *M*_*t*−1_, contributes to forming the next memory state *M*_*t*_. **p**_gen,*t*_ is further transformed into **g**_V,*t*_ in MEC V, from which the predicted sensory observation **x**_*t*_ is decoded. Formally, the generative model defines:

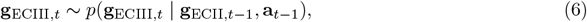

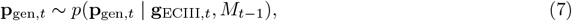

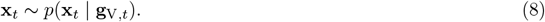

Each conditional distribution is implemented using a fully connected spiking neural network (SNN). Architectural and training details are provided in Algorithm 1 and Table 1.

**Table 1:** Parameters of the model.

| Parameter | Symbol | Value |
| --- | --- | --- |
| Bernoulli sampling steps | $k$ | 2 |
| Number of motor neuron | $\dim(\mathbf{a})$ | 4 |
| Number of CA1 neurons (Inference model) | $\dim(\mathbf{p}_{\mathbf{CA1}})$ | 200 |
| Number of CA1 neurons (Generative model) | $\dim(\mathbf{p}_{\mathbf{gen}})$ | 200 |
| Number of MECII neurons | $\dim(\mathbf{g}_{\mathbf{II}})$ | 200 |
| Number of LECII neurons | $\dim(\mathbf{x}_{\mathbf{LECII}})$ | 200 |
| Number of sensory neurons | $\dim(\mathbf{x})$ | 20 (2D environment), 10 (1D environment) |
| Number of dentate gyrus neurons | $\dim(\mathbf{p}_{\mathbf{DG}})$ | 400 |
| Number of CA3 neurons | $\dim(\mathbf{p}_{\mathbf{CA3}})$ | 200 |
| Number of MECIII neurons | $\dim(\mathbf{g}_{\mathbf{III}})$ | 200 |
| Number of LECIII neurons | $\dim(\mathbf{x}_{\mathbf{LECIII}})$ | 200 |
| Number of MEC V neurons | $\dim(\mathbf{g}_{\mathbf{V}})$ | 200 |
| Action selection interval (steps between decisions) | $T$ | 3 (2D environment), 8 (1D environment) |
| Training iterations | – | 24,000 |
| Batch size | $B$ | 20 |
| Loss weight coefficient | $\lambda_{\text{loss1}}$ | 1.0 |
| Loss weight coefficient | $\lambda_{\text{loss2}}$ | 1.0 |
| Theta modulation input coefficient | $\alpha_{\theta}$ | 10 |
| Threshold voltage | $V_{thr}$ | 0.1 |
| Reset voltage | $V_{reset}$ | -3.0 |
| Minimum membrane potential | $V_{min}$ | -6.0 |
| Maximum membrane potential | $V_{max}$ | 0.9 |
| Membrane time constant | $\tau$ | 0.5 |
| Scaling parameter in surrogate gradient | $\gamma$ | 2.0 |
| STDP potentiation | $A_+$ | 0.0005 |
| STDP depression | $A_-$ | -0.0005 |
| STDP time constant | $\tau_{STDP}$ | 0.5 |
| STDP weight clipping range | $W_{stdp}$ | (-2, 2) |
| Optimizer | – | Adam[26] |
| Weight initialization | – | GlorotUniform[27] |
| Maximum learning rate | $\eta_{max}$ | 0.003 |
| Minimum learning rate | $\eta_{min}$ | 0.0003 |

The memory module *M*, which supports associative storage between the generated pattern **p**_gen,*t*_ and the inferred pattern **p**_CA1,*t*_ during recall, is implemented through synapses undergoing spike-timing-dependent plasticity (STDP) [15][23], defined as:

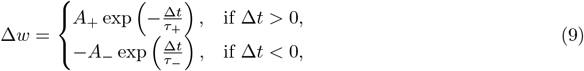

where *A*_+_ and *A*_−_ are the amplitudes, and *τ*_+_ and *τ*_−_ are the time constants for potentiation and depression, respectively. The synaptic change Δ*w* depends on the timing difference Δ*t* = *t*_post_ − *t*_pre_ between presynaptic and postsynaptic spikes.

#### 2.2.2 Inference model

The inference model estimates two latent variables from the current sensory observation **x**_*t*_: (1) the grid cell state **g**_II,*t*_ in MECII, and (2) the place cell state **p**_CA1,*t*_ in CA1 (Fig 1).

CA1 neurons integrate inputs from CA3 and from the lateral entorhinal cortex layer III (LECIII), reflecting its role in combining hippocampal and entorhinal representations. CA3, in turn, forms a recurrent network that combines inputs from the dentate gyrus (DG; **p**_DG,*t*_), MECII state **g**_II,*t*_, and LECII state **x**_LECII,*t*_, together with the previous CA3 state **p**_CA3,*t*−1_. The DG state **p**_DG,*t*_ receives inputs from **g**_II,*t*_ and **x**_LECII,*t*_.

The inference model defines the following probabilistic dependencies:

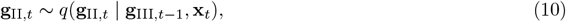

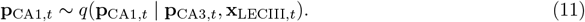

Each conditional distribution is implemented using a fully connected spiking neural network (SNN). Architectural and training details are provided in Algorithm 1 and Table 1.

### 2.2.3 Autoregressive Bernoulli Spike Sampling

In the original Tolman–Eichenbaum Machine (TEM), the grid representation **g** and the place representation **p** were treated as continuous latent variables sampled from Gaussian distributions, enabling differentiable inference in a variational framework. However, in our spiking implementation, both **g** and **p** are modeled as binary spike vectors. To generate these representations in a manner compatible with spiking networks, we adopted the autoregressive Bernoulli

spike sampling method [24], originally developed for fully spiking variational autoencoders. This sampling framework enables the formulation of both the generative model and the inference model using discrete Bernoulli distributions over spike variables. In this method, we first generate pre-activation vaues ***ζ***_*q,t*_ and ***ζ***_*p,t*_ in SNN. These pre-activations are overparameterized: the number of candidate neurons is expanded by a factor of *k* relative to the desired output dimension *C*, resulting in *kC* candidate neurons. These are then divided into *k* non-overlapping groups, each containing *C* neurons (Fig 2). Sampling is performed by randomly selecting one element from each group:

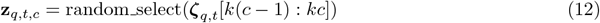

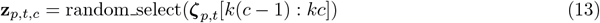

**Figure 2.**
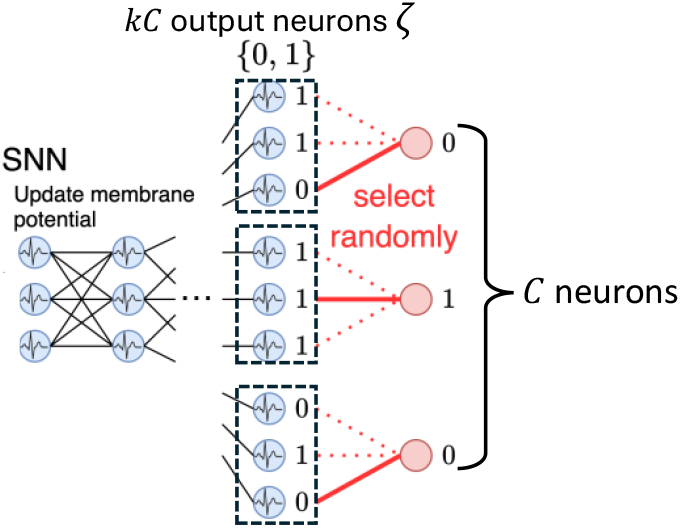
Autoregressive Bernoulli Spike Sampling. This illustrates how sampling from the probability distributions in both the generative and inference models is implemented in a spiking neural network. Example illustration with *k* = 3 and *C* = 3.

By repeating this procedure for 1 ≤*c* ≤*C*, we obtain binary spike vectors **z**_*q,t*_, **z**_*p,t*_ ∈ {0, 1} ^*C*^.

This is equivalent to sampling from the following Bernoulli distributions, where the spike probability for each output neuron *c* is defined as the average of the *k* candidate values in its group:

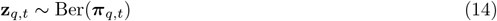

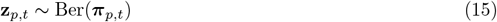

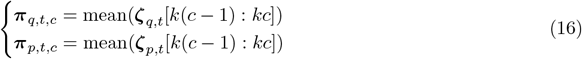

### 2.3 Training procedure

#### 2.3.1 Algorithm

As in TEM, the algorithm is divided into two main phases: (1) Collect data phase, where episodes are executed to store sensory experiences and internal states into a buffer, and (2) Training phase, where the model parameters are updated using sampled batches from the collected episodes. The full procedure is described in Algorithm 1, 2.

##### Algorithm 1 Collect data phase in the SpikingTEM

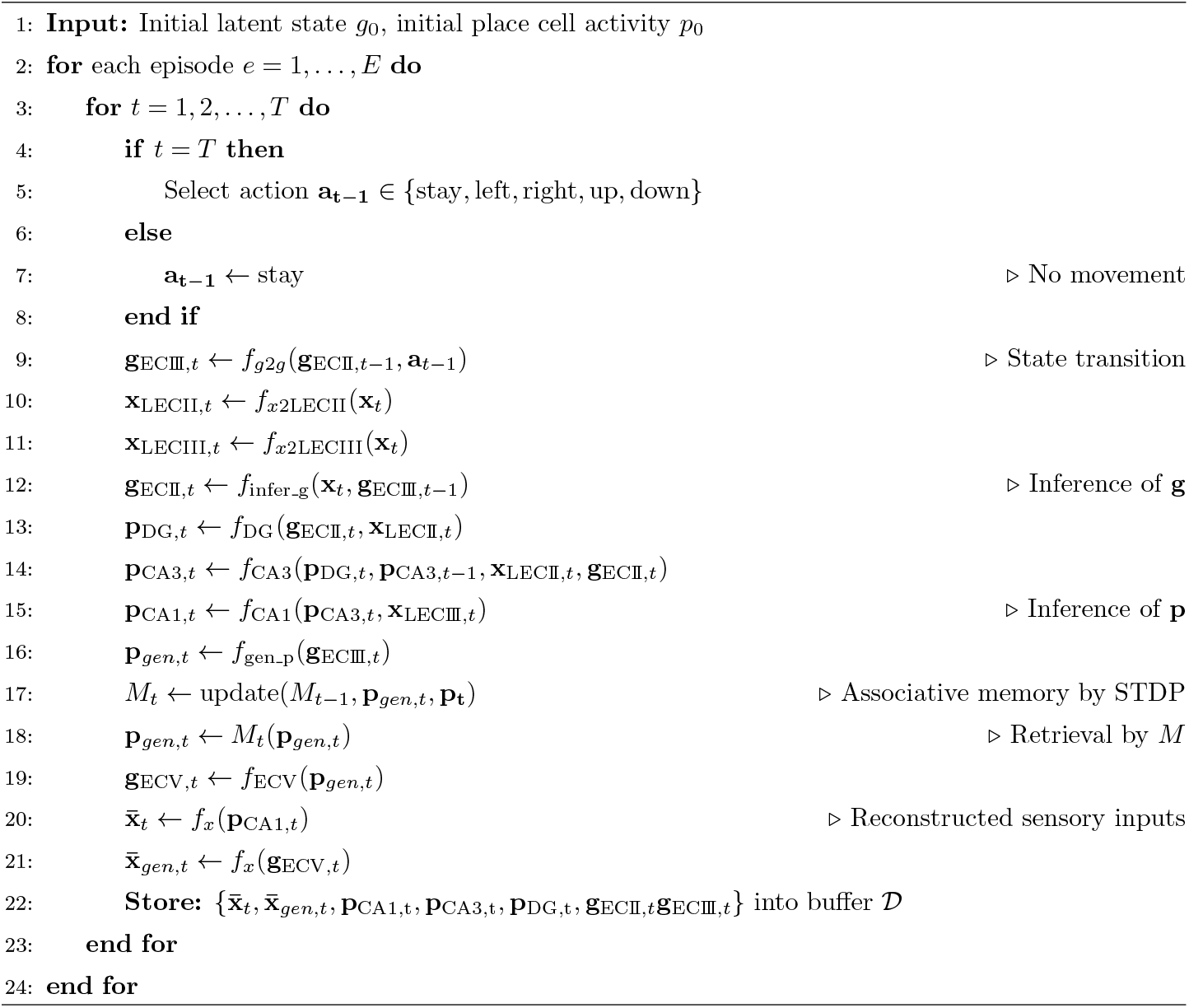

##### Algorithm 2 Training phase in the SpikingTEM

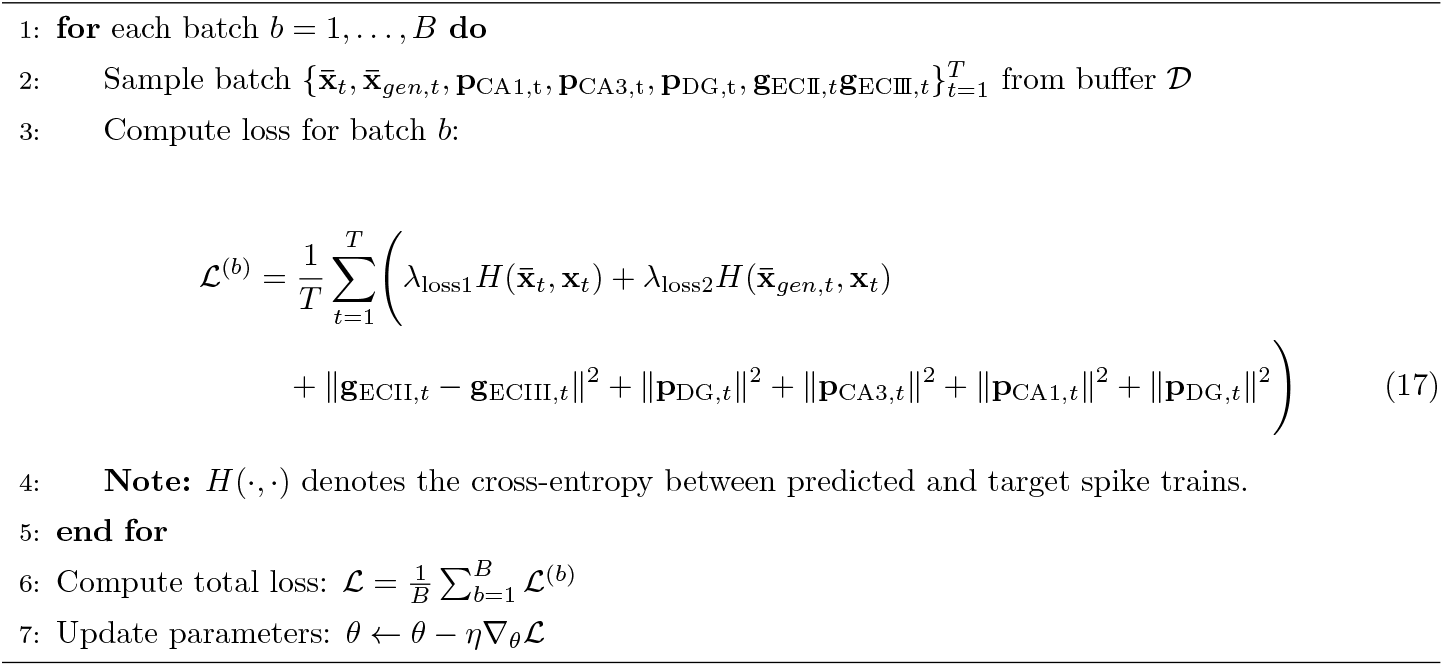

The agent selects a new action (up, down, left, right, or stay) only every *T* steps and remains stationary during the intermediate steps. While it is possible to update the agent s position at every spiking time step, this would require a finer spatial discretization of the environment, resulting in significantly longer simulation times. Therefore, in this work, we simplify the setup by including stay periods between movements.

In the original TEM, the place cell representation **p**_**CA1**_ was computed as a function of the element-wise product of **g** and **x**_**LEC**_, i.e., **p**_CA1,t_ = *f* (**g**_*t*_ ⊙ **x**_LEC,*t*_). However, in the Spiking TEM, using this computation led to the emergence of place cells but failed to produce grid cells (S9 and S10 Figs). Therefore, unlike the original TEM, the element-wise product was not used for the input to **p**_CA1_ in the Spiking TEM.

#### 2.3.2 Surrogate gradient for backpropagation in SpikingTEM

Similar to the original Tolman–Eichenbaum Machine (TEM), we train the SpikingTEM using backpropagation to optimize model weights. However, unlike the original TEM where all computations are differentiable, the use of spiking neurons in SpikingTEM introduces non-differentiable operations due to the discrete nature of spikes. To overcome this challenge, we adopt the surrogate gradient method [25], a standard approach in training spiking neural networks. Although the spiking function is non-differentiable, we approximate its gradient using a smooth surrogate during the backward pass, enabling gradient-based optimization. Let the spike generation be defined as: *s*(*v*) = *H*(*v - v*_*θ*_) where *H*(*·*) is the Heaviside step function and *v*_*θ*_ is the firing threshold. Since 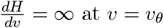, direct backpropagation through *s*(*v*) is not possible. Instead, during training, we use a surrogate derivative: 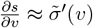 where 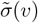 is a smooth function:

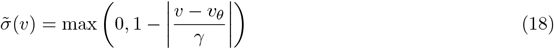

where *γ* is a scaling parameter controlling how broadly the surrogate gradient is spread around the threshold.

### 2.4 Parameter

We summarize the key parameters used in the results (Table 1).

## 3 Results

### 3.1 Spatial navigation in a 2D environment

#### 3.1.1 Emergence of grid cells and place cells

To evaluate whether the Spiking Tolman–Eichenbaum Machine (SpikingTEM) can replicate characteristic spatial firing patterns observed in the hippocampal–entorhinal circuit, we analyzed the neural activities in the entorhinal and hippocampal modules after unsupervised learning in a spatial navigation task.

We trained the model in an 8 *×* 8 square arena, where a virtual agent performed random exploration. At each timestep, the agent selected one of five discrete actions: up, down, left, right, or stay. The number of episodes is 10,000, corresponding to a sequence of movements across the environment. Before the data collection phase, the one-hot observation vectors for each position were randomly assigned to the sensory neurons. These assignments remained fixed throughout each episode, so the agent received the same observation for a given location whenever it visited. No contextual inputs or reward signals were provided; learning was entirely unsupervised. Spiking activity was simulated for *T* time bins per step, and learning proceeded in an unsupervised manner via backpropagation and STDP. The model was trained according to Algorithm 2, with main hyperparameters listed in Table 1.

After training, we evaluated the spatial tuning properties of neurons. To this end, the agent performed an additional 2,000 steps of a random walk in the same 8 *×* 8 environment, without any further learning. In this evaluation phase, the one-hot observation vectors were randomly reassigned to the sensory neurons. This ensures that the test data consisted of novel input patterns, independent of the training data. The spiking activity of neurons during this evaluation phase was recorded and used to compute the spatial firing rate maps. The spatial firing fields and the autocorrelation maps in the medial entorhinal cortex (MECII) module are shown in Fig 3A, B. Fig 3C shows the relationship between the number of SNN training iterations and the proportion of grid cells— neurons in the MECII module with a gridness score [28] greater than 0.8 (See S1 Fig for details). At the beginning of training, no neurons exhibited grid-like activity; however, the number of grid cells gradually increased as learning progressed. Ultimately, grid cells emerged with a 59.6% *±* 18.0% (mean *±* SD). Fig 3D shows the distribution of gridness before and after learning. After training, a larger number of neurons exhibited high gridness values.

**Figure 3.**
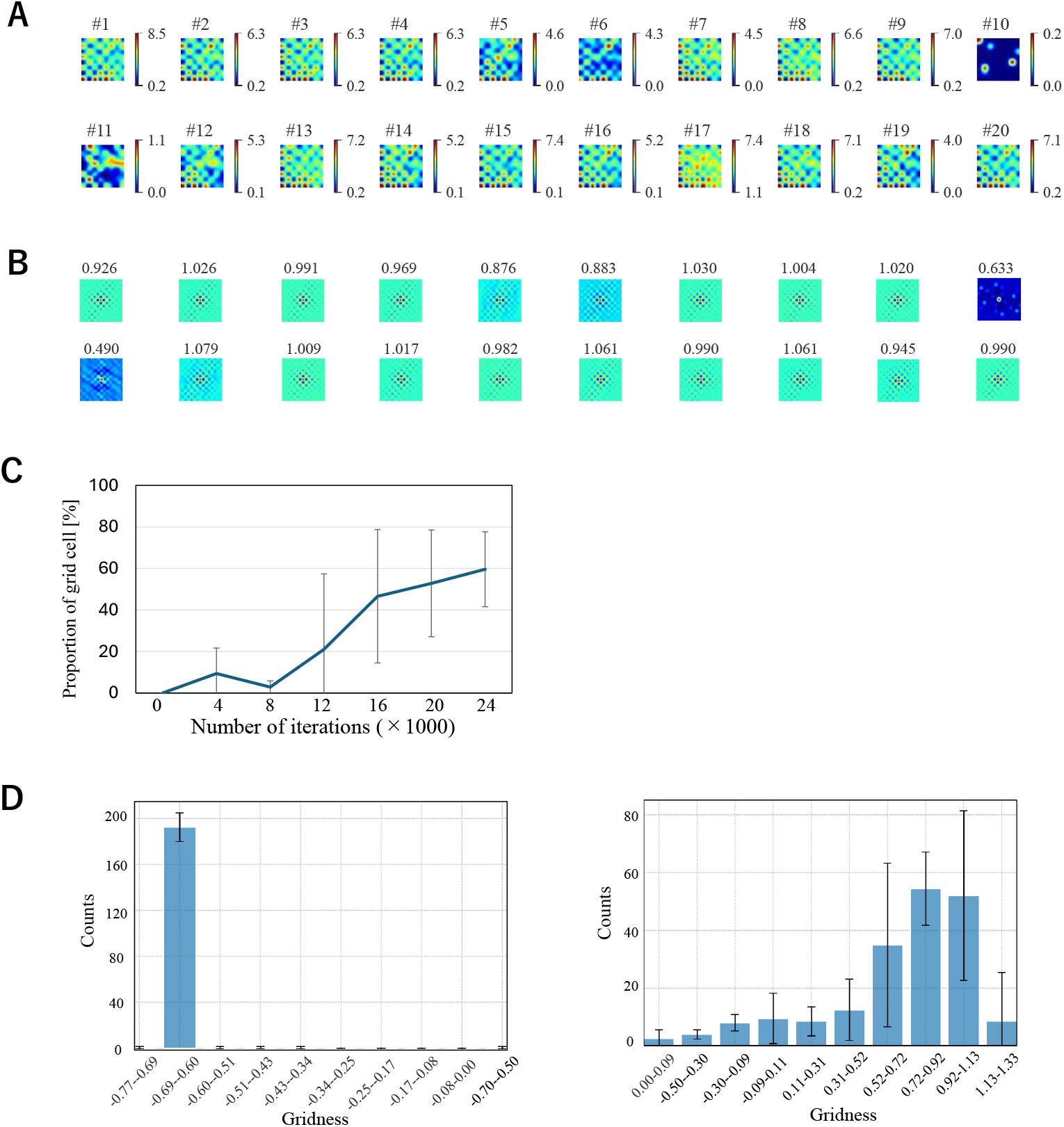
Emergence of grid cells through learning. (A) Spatial firing rate maps of 20 representative MECII neurons after training. Firing rates were computed by averaging neuronal spiking activity over 2,000 steps of random exploration. Spatial firing rate maps for all 200 MECII neurons are shown in S2 Fig. (B) Autocorrelation maps of the same neurons, with the gridness score indicated above each pane. (C) Proportion of grid cells (gridness *>* 0.8) in MECII as a function of training iterations across four independent runs. (D) Distribution of gridness scores before (left) and after (right) training across four independent runs.

The spatial firing fields in the hippocampal module (CA1) after training are shown in Fig 4A. Several neurons exhibit localized, place-specific firing patterns. To quantify this, we computed the spatial information [29] for each neuron, which measures how selectively a neuron fires at specific locations; higher values indicate stronger spatial tuning.

**Figure 4.**
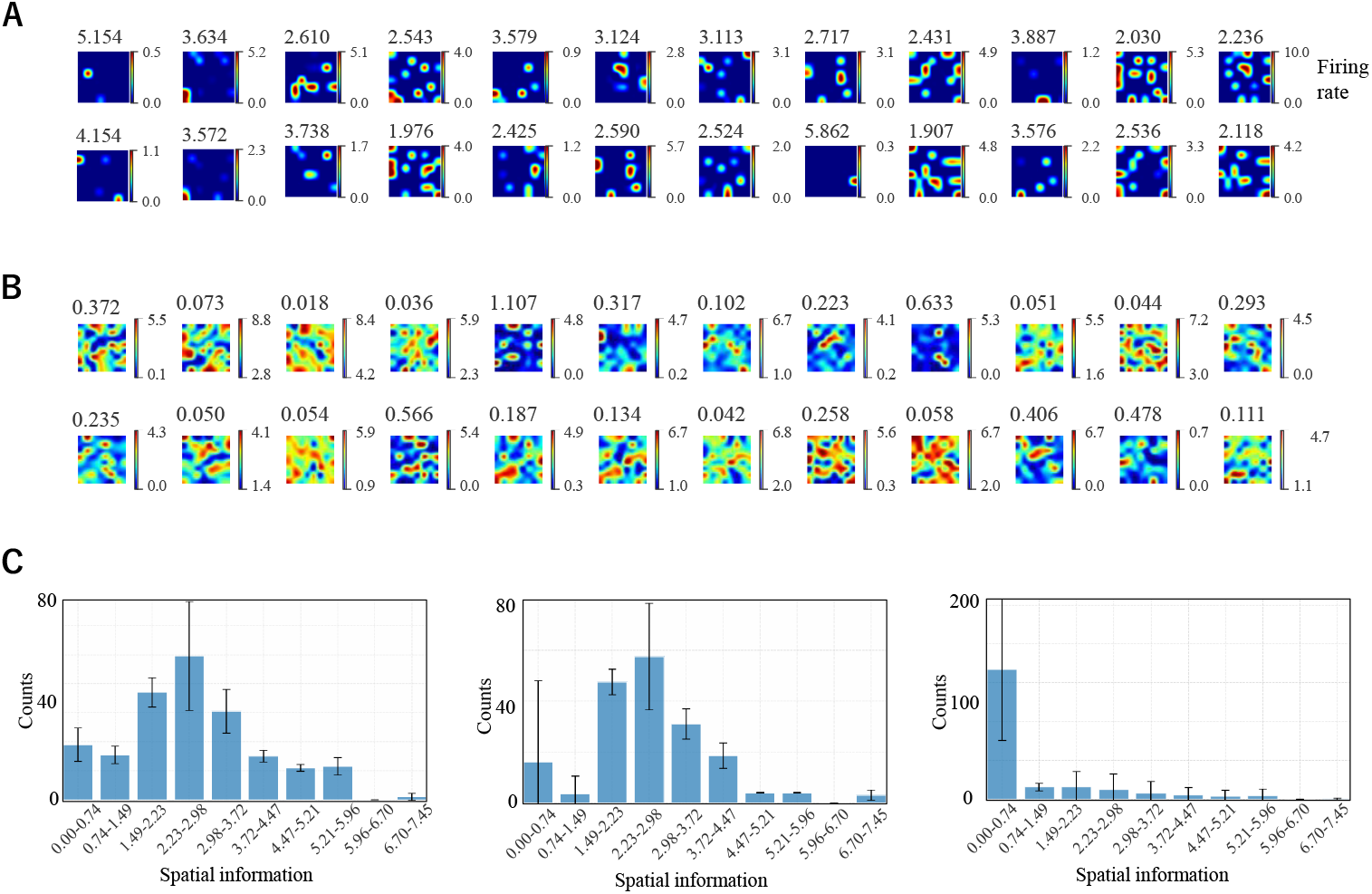
Place cell-like activity in the CA1 module. (A) Example spatial firing rate maps of 24 CA1 neurons after training. The value shown above each map indicates the spatial information score. Firing rates were computed from spiking activity during 2,000 steps of random exploration in the environment. (B) Example firing rate maps of CA1 neurons before training when theta inhibitory modulation was removed. (C) Distributions of spatial information across CA1 neurons under three conditions: before training (left), after training (middle), and before training without theta inhibitory modulation (right). Each results are shown across four independent experiments.

Fig 4B shows the distribution of spatial information values before and after SNN training. Unlike the case of grid cells, there was no substantial change in the distribution of spatial information in CA1 neurons following learning. However, at the beginning of the learning phase, when the theta-modulated inhibitory inputs were removed, most neurons exhibited low spatial information (Fig 4B, C).

In summary, these results demonstrate that grid-like spatial representations spontaneously emerge in the entorhinal cortex through learning, whereas hippocampal place representations depend on the temporal coordination provided by theta-modulated inhibitory inputs.

#### 3.1.2 Realignment and Remapping

Experimental studies have shown that neurons in the hippocampal–entorhinal system reorganize their spatial representations in response to contextual changes. Specifically, grid cells in the entorhinal cortex exhibited global realignment of their grid-like firing patterns across contexts. That is, the grid fields maintained their periodic structure, but the phase and orientation shifted between input conditions [30] — while hippocampal place cells undergo global remapping, where their place fields changed location, disappeared, or re-emerged in different

contexts [31][32]. To test whether our model reproduces this context-dependent spatial coding, we altered the input observation vectors from one-hot to two-hot vectors after training, while keeping the environment ‘s geometry and model parameters fixed. The two-hot input vectors were randomly assigned to sensory neurons for each position in the 8 *×* 8 environment.

Under this condition, neurons in the entorhinal module preserved grid-like periodicity but exhibited realignment across contexts, with their spatial firing fields globally shifting in phase (see Fig 5A, B). Next, we examined the changes in gridness of MECII neurons before and after altering the environment. Despite the altered sensory inputs, the gridness of individual neurons remained largely stable across conditions (Fig 5C). To test the robustness of this effect across different synaptic configurations, we repeated the experiment with two independent models, each having different initial synaptic weights, yielding linear regression fits with *p* = 1.29×10^−5^ and *p* = 2.58 × 10^−18^, respectively, confirming the consistency of gridness preservation across environments.

**Figure 5.**
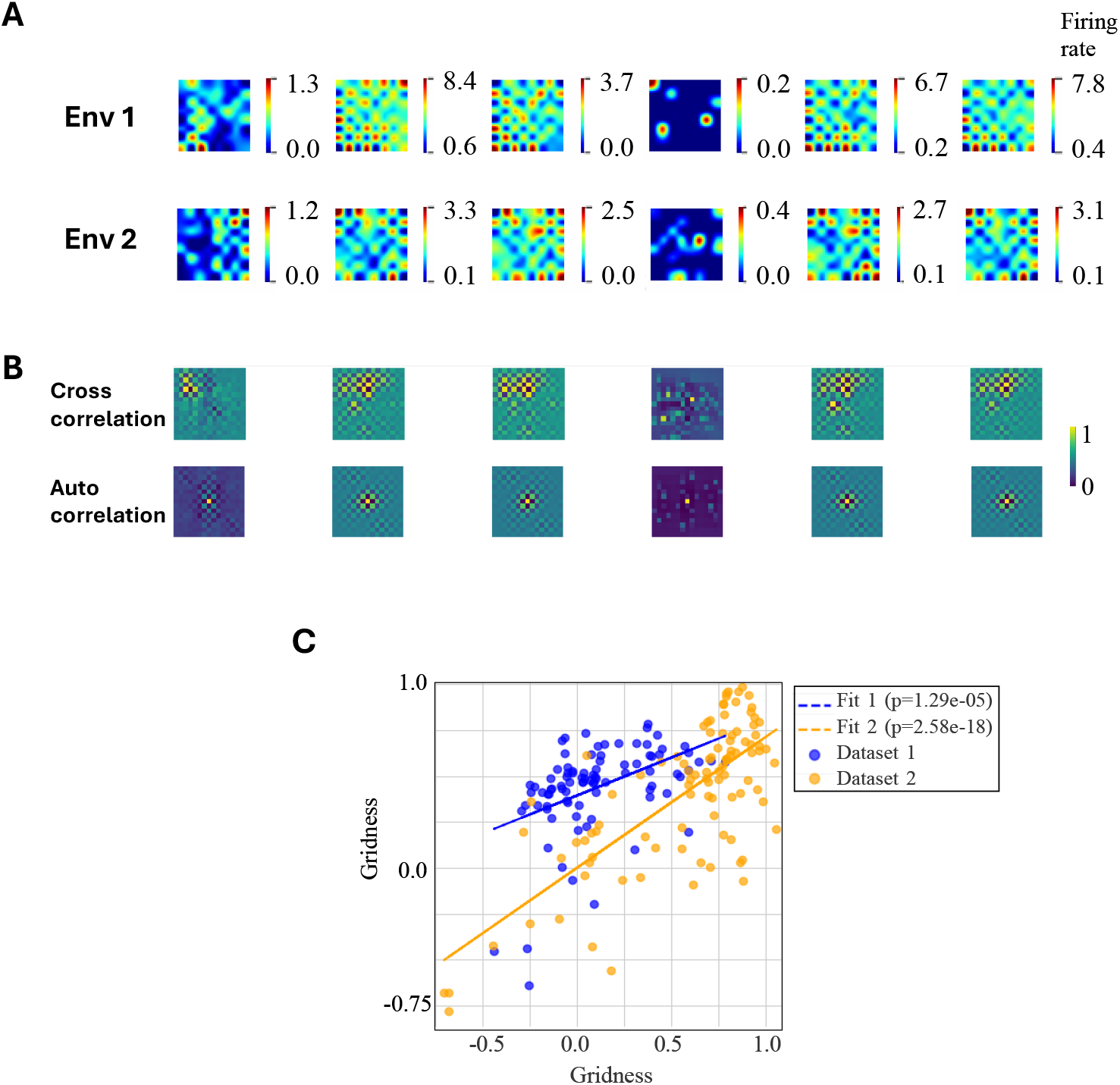
Realignment of grid cell spatial firing patterns in the MECII module across different environments. (A) Examples of firing rate maps from the same six neurons in the MECII module across two environments with different sensory inputs. (B) Cross-correlation maps and autocorrelation maps of the firing rate maps shown in (A). After realignment, the peak positions in the cross-correlation maps are shifted away from the center. (C) Correlation of gridness scores across the two environments for all MECII neurons. Results are shown for two independently trained models with different random weight initializations (Dataset 1 and Dataset 2).

In contrast, neurons in the hippocampal module exhibited global remapping, where place fields relocated under the new input condition (Fig 6). We repeated the experiment with two independent model initializations (i.e., different initial weights), yielding linear regression fits with *p* = 9.17 *×* 10^−35^ and *p* = 4.72 *×* 10^−22^, respectively. This indicates that place-specific firing remained stable across environments (Fig 6B).

**Figure 6.**
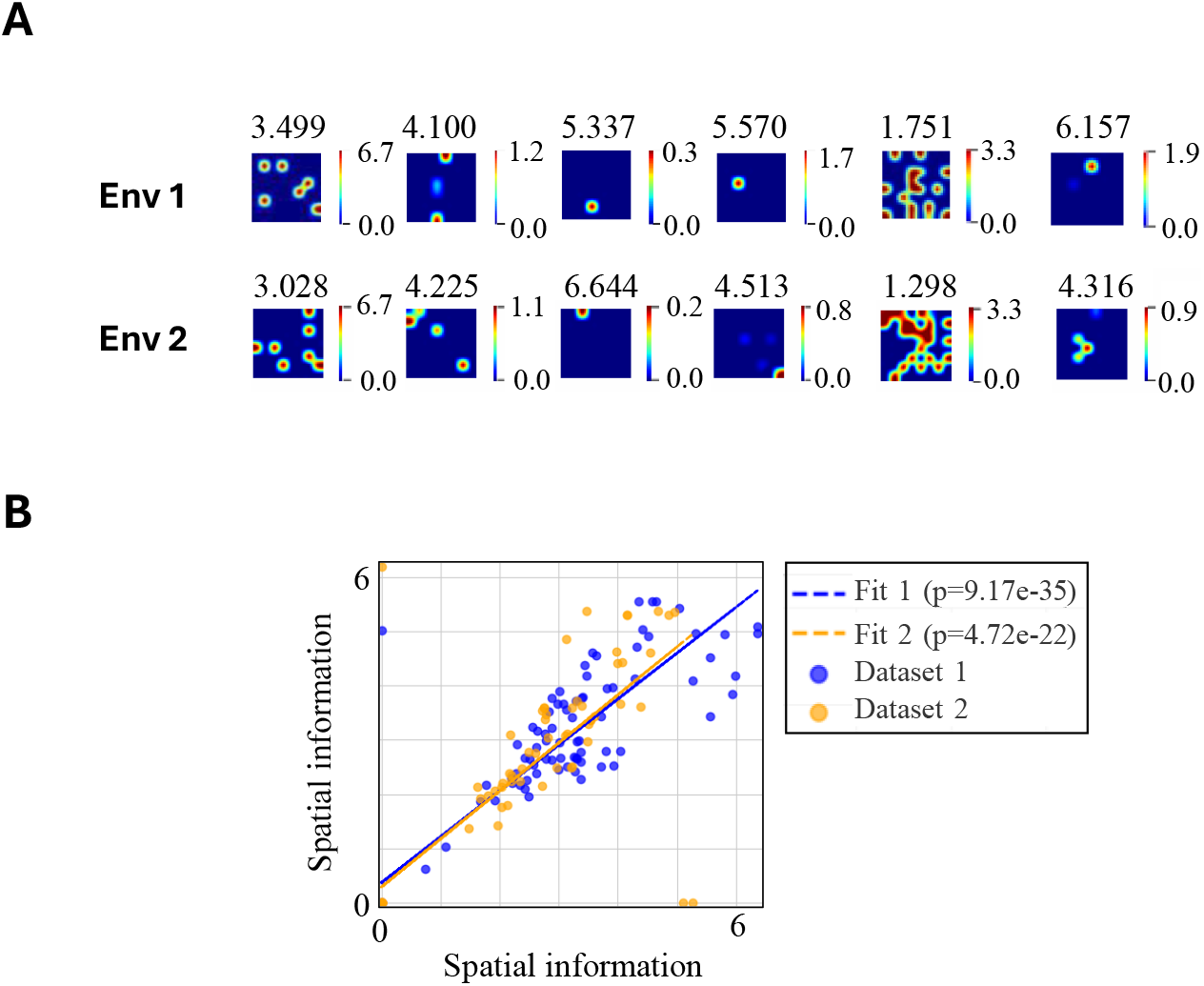
Remapping of spatial firing patterns in CA1 neurons across different environments. (A) Examples of firing rate maps from the same six CA1 neurons recorded in two different environments. The values above each map indicate the spatial information. (B) Correlation of spatial information values across all CA1 neurons between the two environments.

#### 3.1.3 Effect of Plasticity and Circuit Mechanisms on Grid Cell Emergence

To examine the contribution of different mechanisms to the emergence of grid cells, we systematically removed each component of the model and compared the proportion of grid cells in MECII to the baseline condition. As shown in Table 2, the baseline model with all mechanisms active produced the highest proportion of grid cells (59.6% *±* 18.0). Without theta inhibitory modulation *I*_*θ*_ in Eq. 5, the proportion of grid cells was reduced (25.6% *±* 20.6), consistent with previous experimental findings showing that disrupting theta oscillations impairs grid cell formation [16].

**Table 2:**
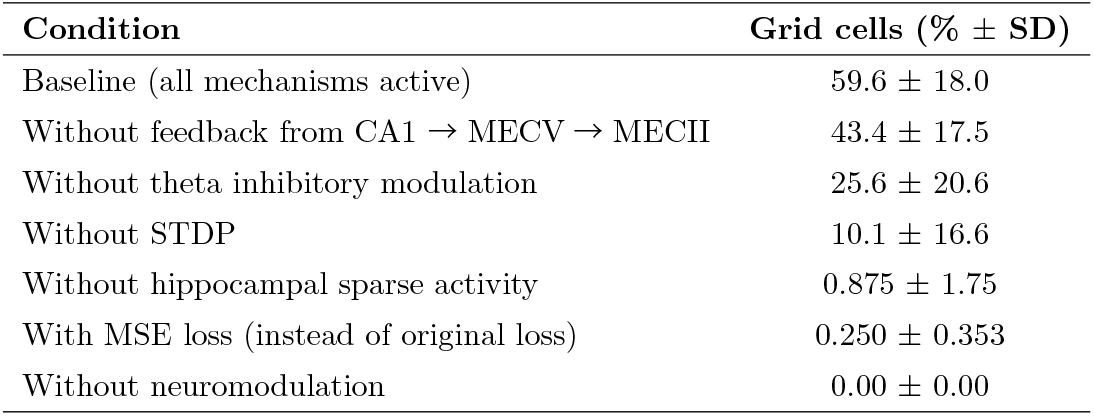
Proportion of grid cells in MECII under different model conditions. Values are mean *±* standard deviation. For each condition, we conducted four simulation runs.

| Condition | Grid cells (% ± SD) |
| --- | --- |
| Baseline (all mechanisms active) | $59.6 \pm 18.0$ |
| Without feedback from CA1 → MECV → MECII | $43.4 \pm 17.5$ |
| Without theta inhibitory modulation | $25.6 \pm 20.6$ |
| Without STDP | $10.1 \pm 16.6$ |
| Without hippocampal sparse activity | $0.875 \pm 1.75$ |
| With MSE loss (instead of original loss) | $0.250 \pm 0.353$ |
| Without neuromodulation | $0.00 \pm 0.00$ |

Removing spike-timing-dependent plasticity (STDP) reduced the proportion of grid cells (10.1% *±* 16.6), indicating that associative memory formation through STDP is essential for the emergence of grid-like representations.

Similarly, removing neuromodulatory signals *G* in Eq. 4 or hippocampal sparsity substantially decreased the proportion of grid cells (0.00% *±* 0.00 and 0.875% *±* 1.75, respectively). Here, hippocampal sparsity refers to the constraint term in the loss function that enforces sparse activity of hippocampal neurons, which in turn promotes structured inputs from the medial entorhinal cortex (MECII) to the dentate gyrus (DG). This finding is consistent with previous models indicating that the sparse activity plays a crucial role in the emergence of grid cells [33]. The reduction in grid cells when theta inhibitory modulation was removed can be explained by its effect on hippocampal activity. Specifically, removing theta inhibitory modulation decreases the spatial information of hippocampal neurons (Fig 4B) and reduces the sparsity of hippocampal firing. This, in turn, diminishes the ability of the network to generate robust grid-like patterns in the entorhinal module.

Finally, replacing the original objective function with a simple mean squared error (MSE) loss also impaired grid cell formation (0.250% *±* 0.353), suggesting that the loss function plays an essential role in stabilizing spatial representations.

Notably, removing feedback from CA1 to MECV →MECII reduced the proportion of grid cells to 43.4% *±* 17.5, but this difference from the baseline was not statistically significant (p=0.244) Together, these results demonstrate that theta inhibitory modulation, STDP, neuromodulation, and hippocampal sparse activity are all necessary for the robust emergence of grid cells.

#### 3.1.4 Effect of Sensory Ambiguity on Grid Cell Formation

We investigated how the uncertainty between sensory inputs and spatial locations influences the emergence of grid cells. In this study, neurons with a gridness score greater than 0.8 were classified as grid cells.

In our setup, each position in the 8 *×* 8 arena was associated with a one-hot sensory input, where one of the sensory neurons was active for each location. When the total number of sensory neurons matched the number of positions (8 *×* 8 = 64), there was a one-to-one correspondence between sensory input and spatial location, and the position could be uniquely inferred from sensory input alone. However, when the number of sensory neurons was reduced, this correspondence became ambiguous —multiple positions could produce the same sensory input —introducing uncertainty in spatial representation.

As shown in Fig 7, grid cells emerged most robustly when this ambiguity was moderate (around 20 sensory neurons). When the number of sensory neurons was either too small or too large, the proportion of neurons classified as grid cells decreased sharply. These results suggest that grid-like internal representations emerge as a compensatory mechanism under conditions where sensory information alone is insufficient to uniquely determine spatial location (see Discussion for details).

**Figure 7.**
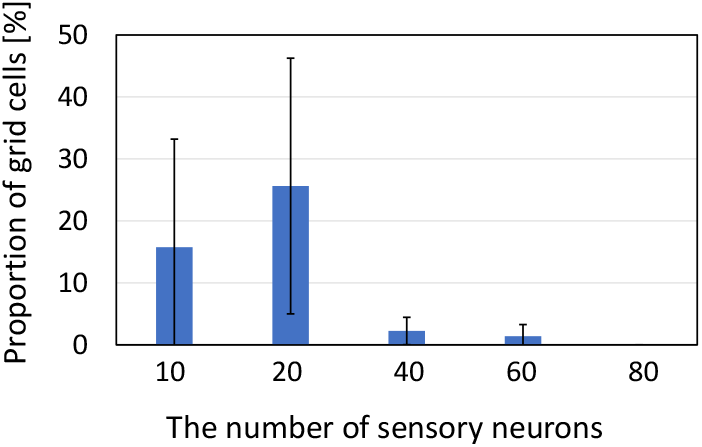
Effect of sensory population size on grid cell emergence. The proportion of grid cells in MECII is plotted against the number of sensory neurons. Each condition was tested in four experiments.

#### 3.1.5 Predictive Grid Cells

In our model, neurons in the MECIII module were trained to predict the activity of MECII neurons one step ahead in time. Interestingly, the spatial firing fields of MECIII neurons based on the current position exhibited relatively low gridness. However, when the activity was evaluated using the position at the next time step (one step ahead), the gridness of these neurons increased substantially (*p* = 2.983 *×* 10^−6^)(Fig 8). This indicates the presence of *predictive grid cells* in MECIII, whose spatial tuning anticipates future positions. These results are consistent with experimental findings reporting predictive coding properties in MECIII neurons [34].

**Figure 8.**
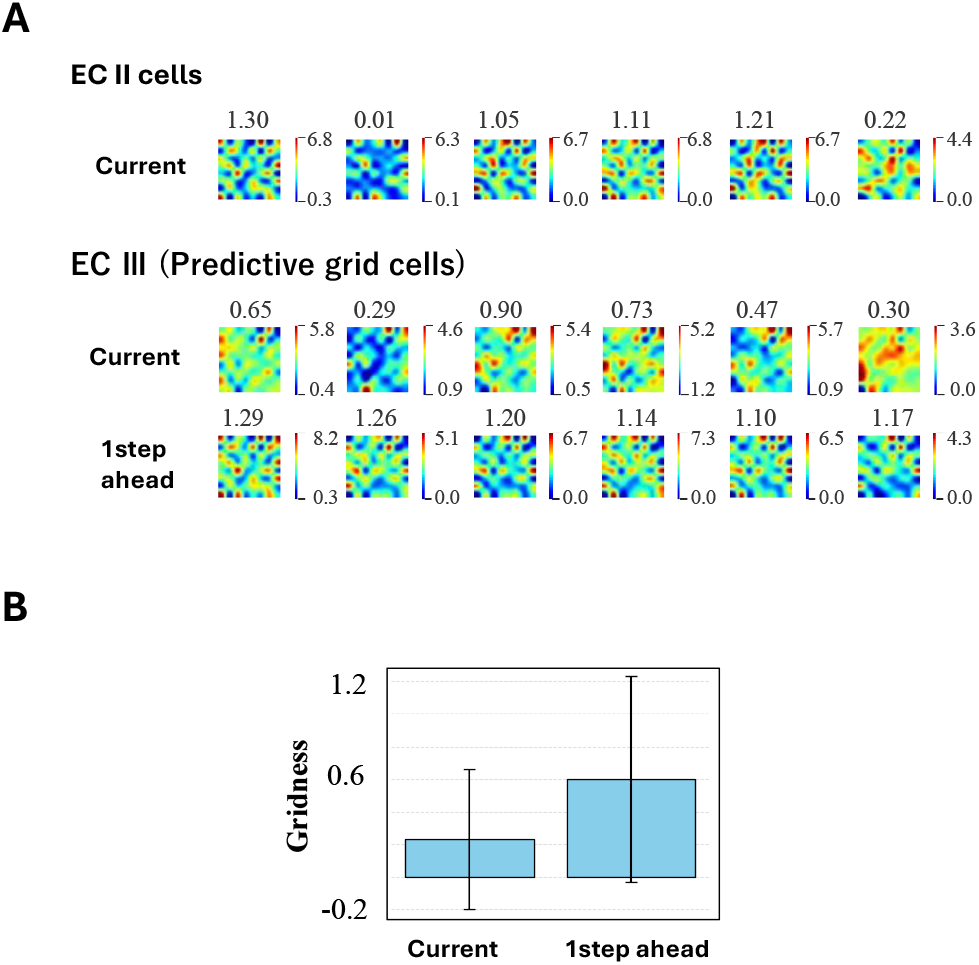
Predictive grid cells in MECIII. (A) Examples of firing rate maps for six neurons in the MECII and MECIII modules after training. Values above each map indicate the gridness score. For MECIII neurons, computing the firing rate map using the position one step ahead results in higher gridness values. Spatial firing rate maps for all 200 MECIII neurons are shown in S3 Fig. (B) Gridness scores for all MECIII neurons computed using the current position versus the position one step ahead (*p* = 2.983 *×* 10^−6^).

### 3.2 Spatial navigation in a linear track

To evaluate whether the model forms spatially selective representations in a linear track setting, we tested it in a one-dimensional environment consisting of 40 discrete positions. The agent moved back and forth along the track for 5,000 episodes, advancing by one position per episode and reversing direction at each end. Each episode was further divided into *T* = 8 temporal bins. Before the data collection phase, the one-hot observation vectors corresponding to each position were randomly assigned to the sensory neurons. These assignments remained fixed throughout all episodes, such that the agent received the same observation whenever it visited the same location. No explicit goals or external cues were provided. The model was trained according to Algorithm 2, as in the two-dimensional environment.

After training, the agent performed an additional 1,000 episodes without any further learning. Before this evaluation phase, the one-hot observation vectors were randomly reassigned to sensory neurons for each position, ensuring that the sensory inputs differed from those used during training. After the evaluation phase, we examined the spatial firing patterns of neurons. Neurons in the entorhinal module developed grid-like firing with periodic activation along the track (Fig 9), while neurons in the hippocampal module formed sharply localized place fields (Fig 10).

**Figure 9.**
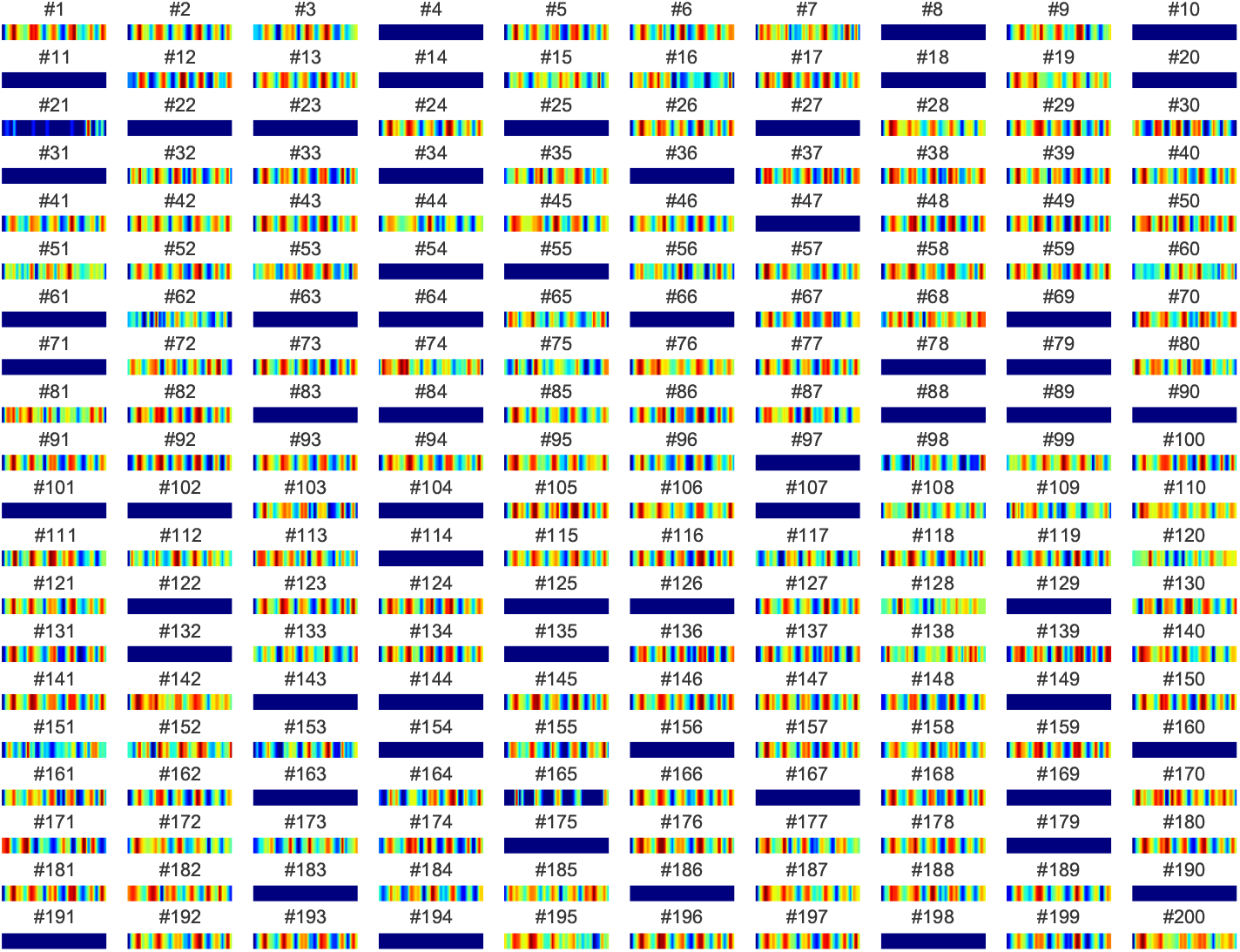
Spatial firing fields of MECII neurons on a linear track. Each panel shows the firing rate map of a neuron in the MECII module recorded during 20 laps of a linear track. Neurons exhibit diverse spatial tuning profiles, including periodic firing fields along the track.

**Figure 10.**
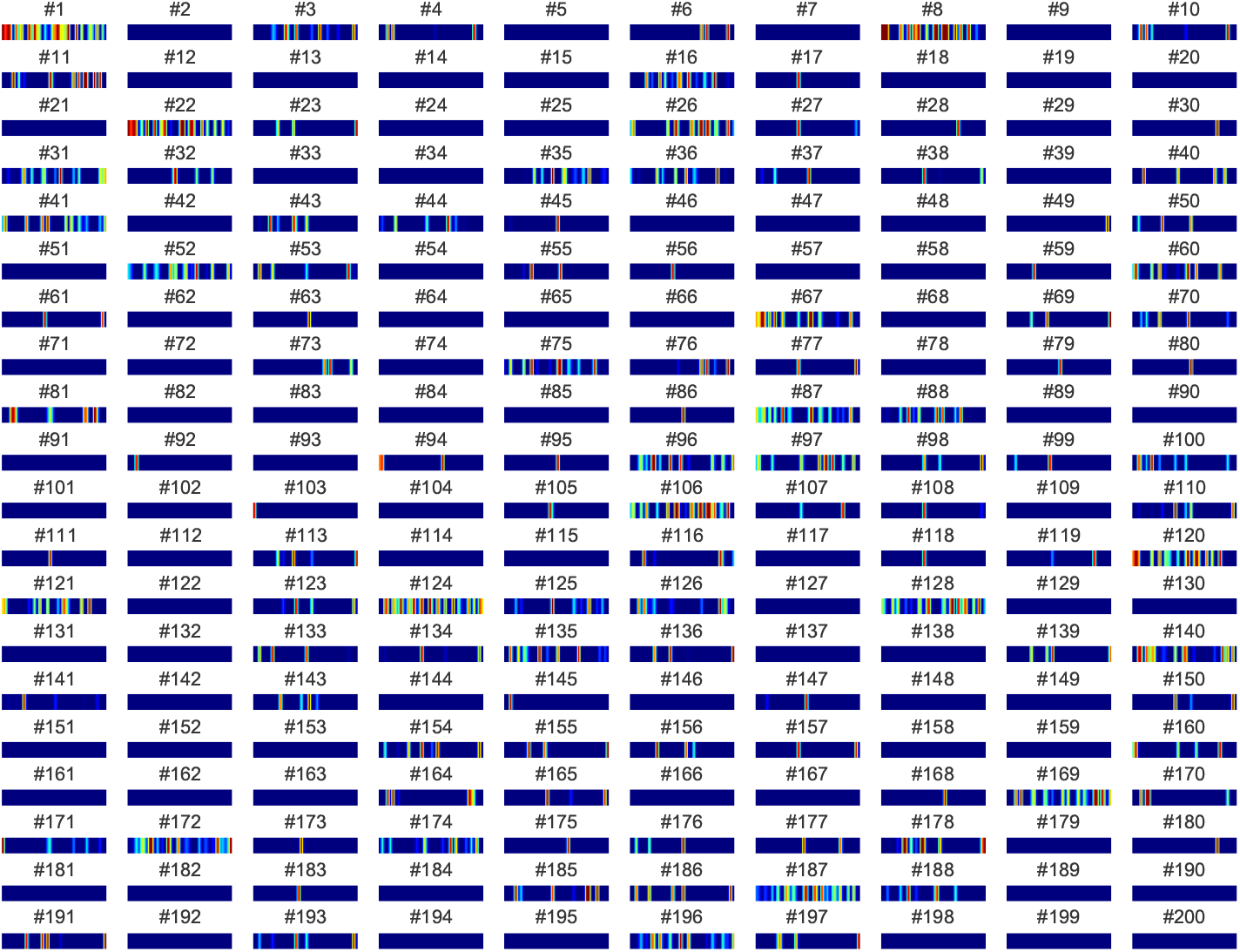
Spatial firing fields of CA1 neurons on a linear track. Each panel shows the firing rate map of a neuron in the CA1 module recorded during 20 laps of a linear track.

#### 3.2.1 Phase precession and phase-locking of grid cells

We next examined the temporal relationship between grid cell firing and the theta rhythm. For this analysis, spikes were aligned to the local theta oscillation, and their phases were plotted as a function of the animal’s position along the linear track. In some grid cells, spikes advanced systematically to earlier theta phases as the animal traversed the firing field, corresponding to phase precession (Fig 11). In other cells, spikes were concentrated around a restricted range of theta phases, regardless of position, indicating phase locking to the ongoing theta oscillation (Fig 12).

**Figure 11.**
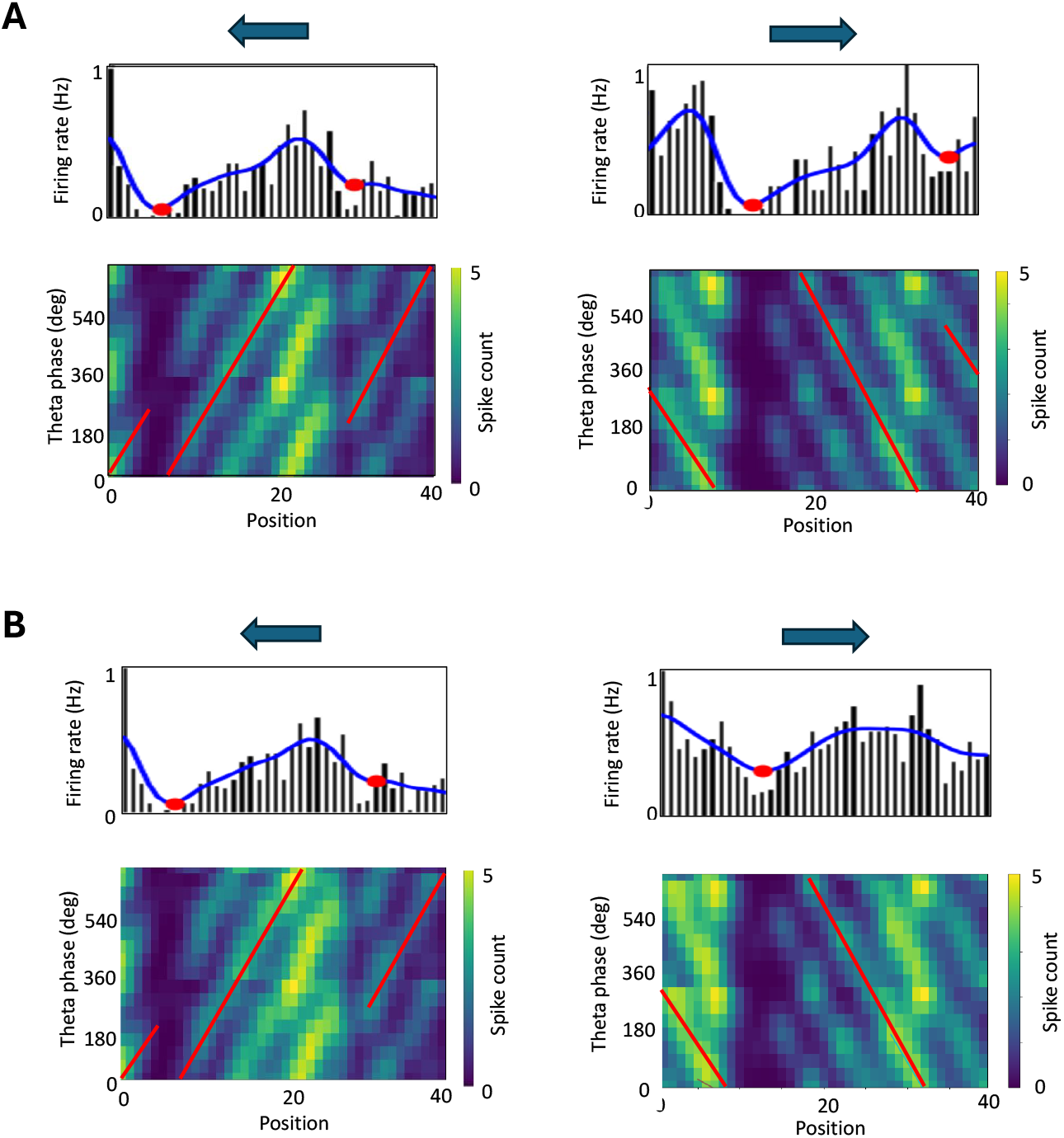
Phase precession examples of MECII and MECIII neurons. (A) Example of phase precession in an MECII neuron. The arrow indicates the running direction along the linear track. The blue line represents the Gaussian-smoothed firing rate, and red dots denote local minima. The lower panel shows the phase–position heatmap. The trajectory was segmented by the local minima, and a circular–linear regression was performed (red line). (B) Example of phase precession in an MECIII neuron. The firing rate and phase–position diagram were computed using the position one step ahead, because MECIII neurons predict the activity of MECII neurons in the next step. A circular–linear regression was performed (red line). Cells were classified based on the absolute slope of the regression line: values ≤ 0.2 [rad/position bin] were defined as *phase-locked*, between 0.2 and 0.6 as *phase-precession*, and ≥ 0.6 as *phase-independent*.

**Figure 12.**
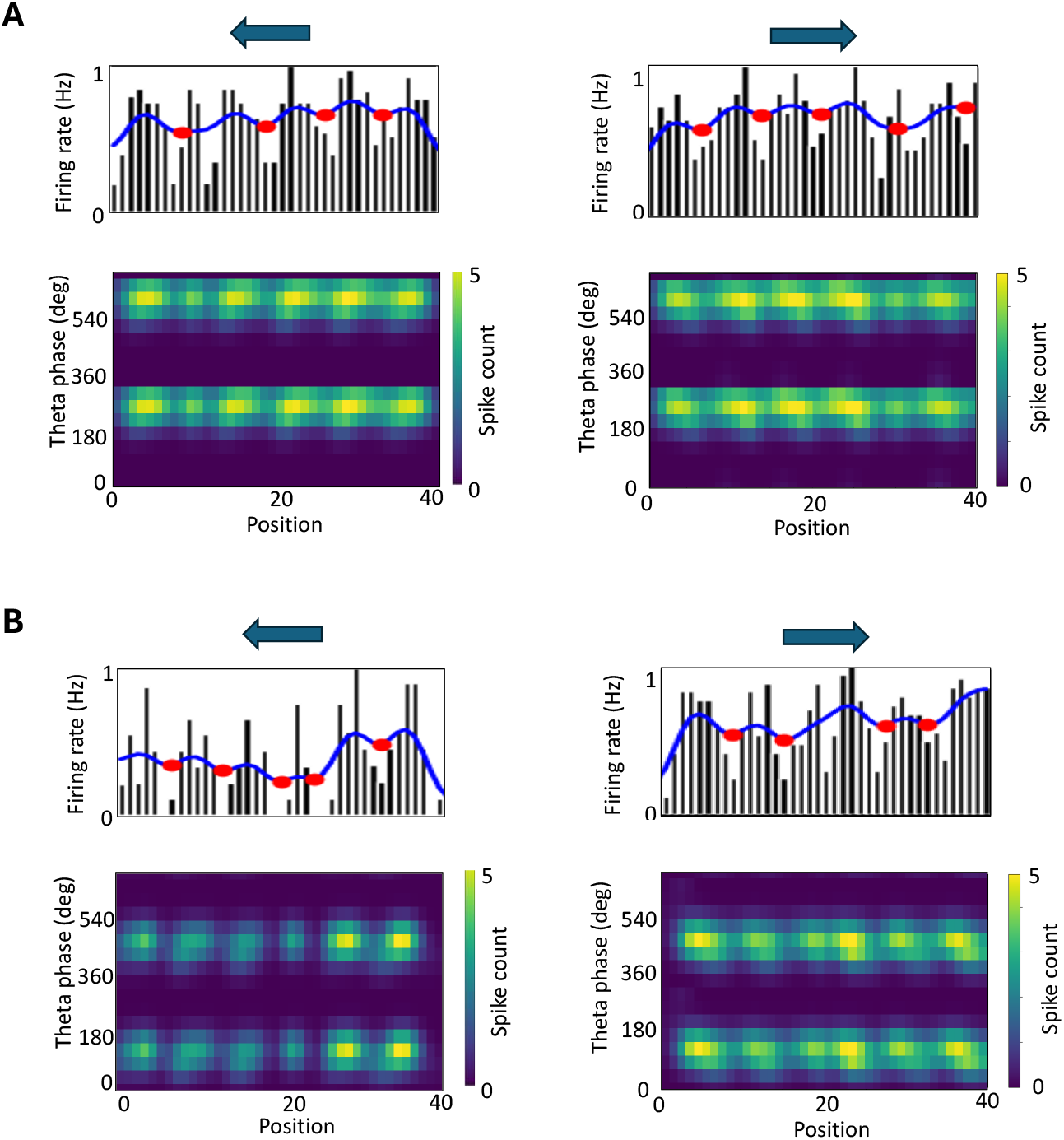
Representative examples of phase locking of grid cell firing along a linear track. Example MECII neuron. The top panels show the firing rate of the cell as a function of position for the forward (left) and backward (right) runs. The lower panel shows the phase–position diagram. This example exhibits clear phase locking, with spikes consistently occurring at a specific theta phase. (B) Example MECIII neuron. Similar to (A), the neuron shows phase locking.

We next examined how grid cells in MECII and MECIII exhibited phase precession or phase locking under different modulatory conditions. When neuromodulation *G* — which controls the gain of input current (Eq. 4) was enabled and theta inhibitory modulation *I*_*θ*_ (Eq. 5) in MECIII was disabled, all grid cells in both MECII and MECIII exhibited phase precession (Fig 13A). In contrast, when neuromodulation was disabled and theta inhibitory modulation in MECIII was enabled, all grid cells in both regions exhibited phase locking (Fig 13B). Finally, when both neuromodulation and theta inhibitory modulation in MECIII were enabled, a mixed pattern emerged: in MECII, the majority of grid cells showed phase precession (80.2 ± 19.3%), whereas in MECIII, most grid cells exhibited phase locking (86.7 ± 11.5%) (Fig 13C). These results are consistent with experimental findings showing that grid cells in MECII exhibit clear phase precession relative to the theta rhythm, whereas neurons in MECIII tend to be phase-locked to a specific theta phase [11][35].

**Figure 13.**
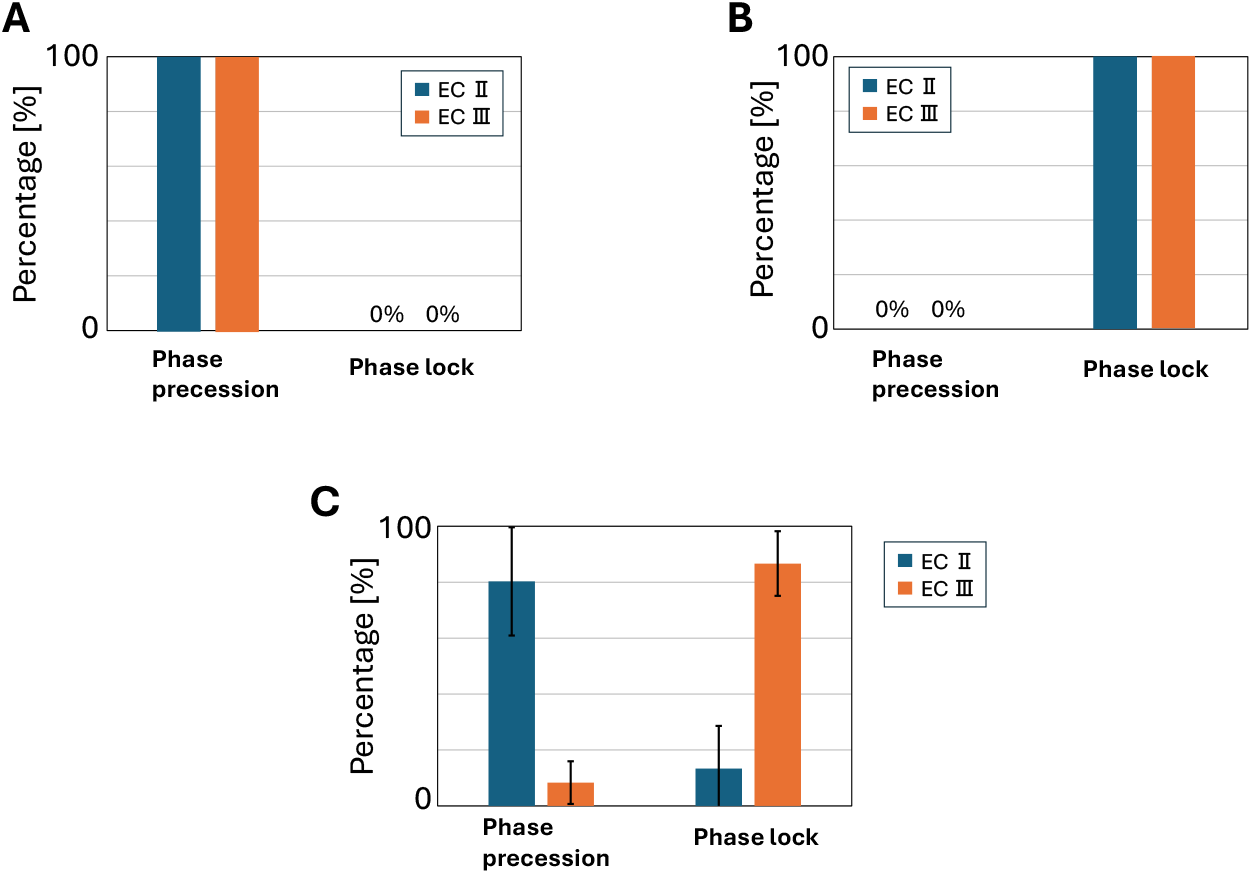
Relationship between neuromodulation, theta inhibitory modulation, and phase precession/phase locking. (A) Percentages of phase-precessing and phase-locked grid cells in MECII and MECIII when neuromodulation was enabled and theta inhibitory modulation in MECIII was disabled. (B) Percentages of phase-precessing and phase-locked grid cells in MECII and MECIII when neuromodulation was disabled and theta inhibitory modulation in MECIII was enabled. (C) Percentages of phase-precessing and phase-locked grid cells in MECII and MECIII when both neuromodulation and theta inhibitory modulation in MECIII were enabled.

These results suggest that neuromodulation promotes the temporal advancement of spikes (phase precession), while theta inhibitory modulation stabilizes spike timing (phase locking), and that the interplay between these mechanisms shapes distinct temporal coding dynamics across MECII and MECIII.

### 3.3 Analysis of synaptic inputs to grid cells and place cells

To gain insights into the mechanisms underlying spatial selectivity, we analyzed the synaptic inputs received by grid cells in the entorhinal cortex and place cells in the hippocampus.

#### 3.3.1 Grid Cell Weight Matrices between MECII and MECIII

To examine the interactions between MECII and MECIII in the Spiking TEM, we analyzed the synaptic weight matrices connecting MECII to MECIII under different movement conditions. Five separate matrices were computed, corresponding to the actions: left, right, up, down, and stay.

Fig 14A shows an example of the firing rate maps and weights from MECII to a single MECIII grid cell in MECIII. Fig 14B displays the distribution of weights from MECII to a single MECIII grid cell for each action (left, right, up, down, and stay). Fig 14C shows the average weights to all MECIII grid cells. We examined two models with weights randomly initialized with both positive and negative values (GlorotUniform initialization [27]), and in both cases, after learning, inhibitory weights were dominant regardless of the action. This finding is consistent with experimental observations indicating that grid cell circuits are largely inhibitory [36].

**Figure 14.**
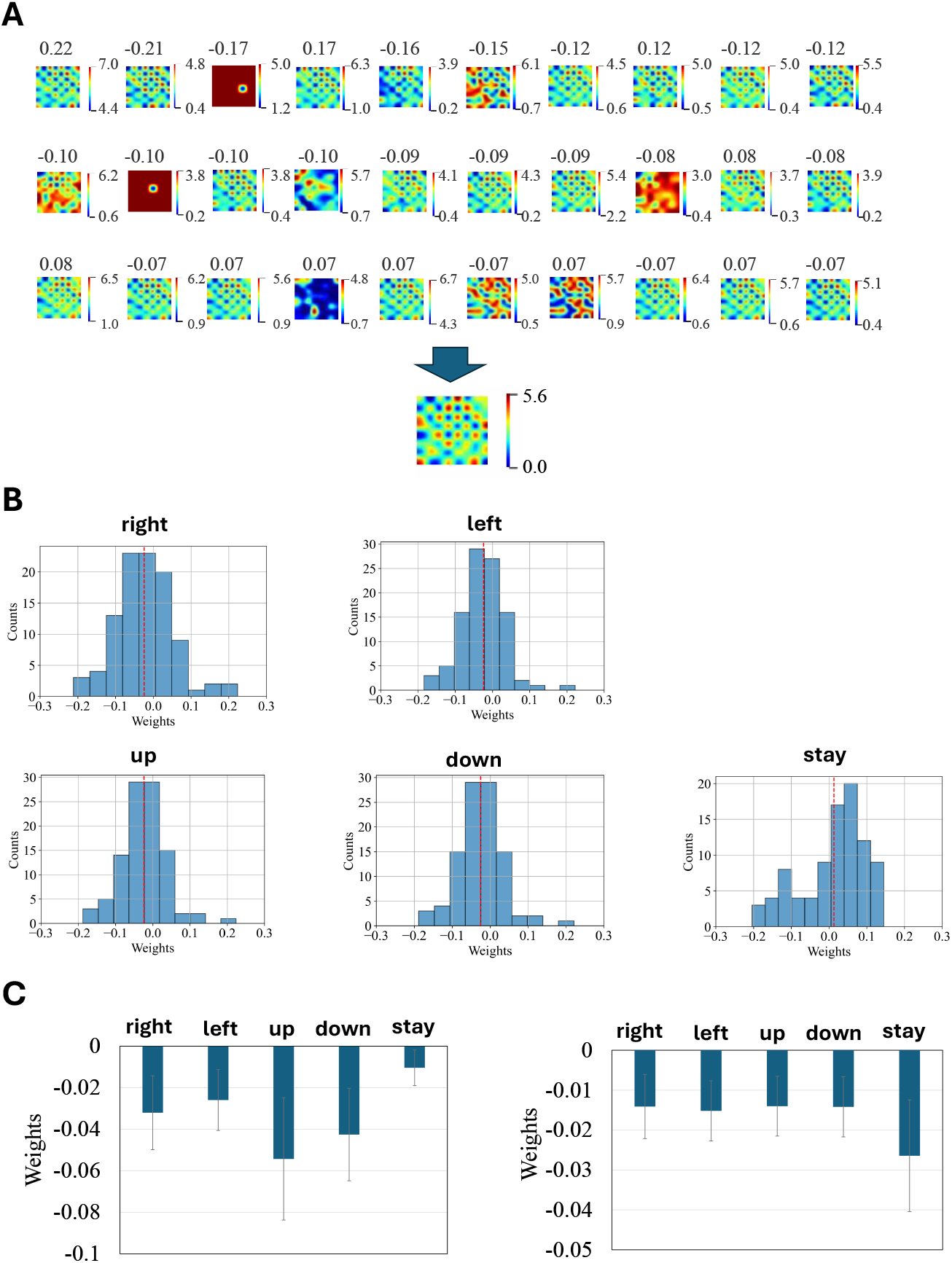
Synaptic inputs to grid cells in MECIII. (A) Each panel shows the spatial firing fields of presynaptic neurons in MECII projecting to a selected grid cell in MECIII. The value above each panel indicates the synaptic weight. Panels are ordered by descending absolute weight. (B) Distribution of MECII → MECIII transition matrix weights for five actions (right, left, up, down, stay). The red line indicates the mean. (C) MECII → MECIII transition matrix weights for each of the five actions after training two models with different initial synaptic weights.

#### 3.3.2 Place Cell Weight Matrices between MECIII and CA1

For place cells, we plotted the weights from the grid module (**g**_**ECIII**_) to a hippocampal neuron (Fig 15). Most of the inputs received by CA1 place cells from MECIII grid cells were inhibitory, with only a small fraction of excitatory connections selectively targeting specific neurons. This sparse and selective pattern of excitatory inputs likely underlies the formation of localized place fields, while the predominant inhibition serves to suppress non-specific activity and enhance spatial selectivity.

**Figure 15.**
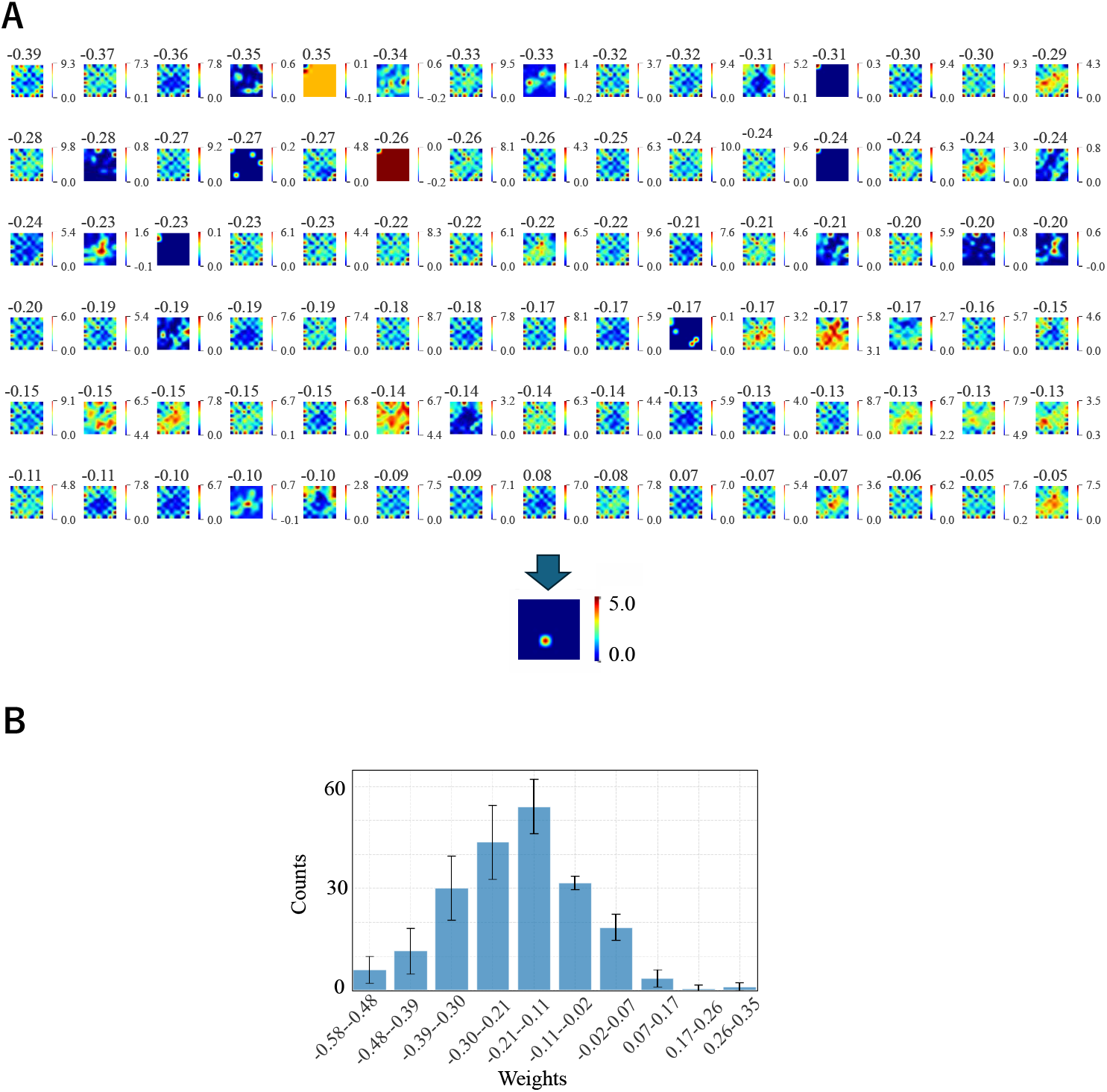
Synaptic inputs to place cells in the CA1 module. (A) Each panel shows the spatial firing fields of presynaptic neurons in MECIII projecting to a selected place cell. The value above each panel indicates the synaptic weight. Panels are ordered by descending absolute weight. (B) Distribution of synaptic weights for CA1 neurons with spatial information greater than 5.

## Discussion

Our spiking neural network implementation of the Tolman–Eichenbaum Machine (SpikingTEM) successfully reproduced key biological phenomena, such as the emergence of grid cells and place cells.

We observed that a large proportion of neurons remained silent throughout the simulation. This is consistent with experimental findings showing that in a given environment, only about 40% of hippocampal pyramidal neurons are active, while the remaining 60% are silent cells [37]. In the Spiking TEM, 302 out of 600 hippocampal neurons (50.3%) were classified as silent cells (S4, S5, S6 Fig). This ratio is comparable to experimental observations. Notably, the dentate gyrus exhibited particularly sparse activity, with 180 out of 200 neurons (90.0%) remaining silent — closely matching the high proportion of silent granule cells reported in vivo (163 out of 190 neurons (85.8%), [38]). Overall, these results demonstrate that the Spiking TEM reproduces the characteristic sparsity of hippocampal activity observed in biological neural circuits.

Furthermore, although with low probability, larger-scale grid patterns occasionally emerged in our simulations (observed in 1 out of 20 runs) (S7, S8 Fig). This observation is consistent with experimental findings showing that grid scale increases along the dorsal–ventral axis of the entorhinal cortex [1]. These results suggest the potential existence of a hierarchical organization of spatial representations, and elucidating the factors that determine grid scale remains an important question for future studies.

With respect to the results shown in Fig 7, the observed dependence of grid cell emergence on the number of sensory neurons can be interpreted in terms of the balance between external sensory inputs and internal spatial representations. When the number of sensory neurons is relatively small compared to the number of environmental states (i.e., the number of spatial bins in the arena), the sensory input alone is insufficient to uniquely determine the animal ‘ s position. Under such conditions, the network benefits from forming an internal code, namely the grid-like representation **g**, which provides a structured basis for disambiguating locations and supporting path integration. This explains why a higher proportion of grid cells was observed when the sensory population size was limited. In contrast, when the number of sensory neurons is large, the sensory input itself becomes sufficiently informative to distinguish positions across the environment. In this case, the reliance on internal representations is reduced, and grid cells are no longer necessary for accurate position estimation. As a result, the proportion of neurons classified as grid cells decreases sharply. These findings suggest that the emergence of grid-like coding depends on the relative information content of external versus internal sources. Grid cells are most prominent when sensory information is sparse or ambiguous, highlighting their role as an internal coordinate system that complements incomplete external cues. Conversely, when external information is abundant, the necessity of such an internal system diminishes, leading to a suppression of grid-like activity.

Despite these promising results, several challenges remain for future work. First, to reduce computational costs, we simulated the network with an enlarged time resolution compared to the actual timescale of biological spikes. Refining the simulation with a finer temporal resolution would allow for a more accurate reproduction of temporal coding phenomena. However, in this study, we employed Backpropagation Through Time (BPTT), which becomes computationally expensive for learning over long temporal intervals. Developing more efficient learning algorithms capable of handling long timescales remains an important challenge. Temporal coding is not limited to phase precession and phase locking, but also includes replay [39]. Recent studies have leveraged replay to implement biologically plausible, rapid one-shot learning versions of BPTT [40]. Additionally, replay has been studied in continuous attractor networks in relation to grid cell formation [41]. A unified model that can account for both temporal and spatial coding, integrating these mechanisms, remains an open challenge for future research.

Second, in our experiments, the emergent grid cells predominantly exhibited square or diamond-shaped grid patterns, whereas canonical hexagonal grids were not observed. Previous modeling studies have reported that constraining the weights from the entorhinal cortex to the hippocampus to be non-negative can lead to a transition from square to hexagonal grid patterns [42][43]. This suggests that weight constraints may play a critical role in shaping the geometry of grid representations. An important direction for future work is therefore to examine whether introducing such non-negativity constraints in our model can give rise to hexagonal grid cells, thereby bridging the gap between our findings and biological observations.

Third, although grid-like firing fields emerged in our model, the diversity of grid cells was limited. In particular, the model failed to reproduce the variability in grid phases — that is, spatial offsets of the grid pattern relative to the environment. This lack of diversity likely prevented the emergence of toroidal structure in the population activity of grid cells, which has been observed in biological data [44]. Addressing this limitation may require revisiting architectural components of the model, such as the network connectivity or learning rules, as well as tuning hyperparameters.

Finally, although we employed backpropagation for learning, it remains controversial whether such a mechanism is used in the brain [45]. More biologically plausible local learning rules— such as predictive coding or biologically realistic implementations of backpropagation [46]—may provide viable alternatives. While grid cells have been shown to emerge in rate-based predictive coding models [47], adapting such mechanisms to a spiking framework is an important step toward bridging theory and biology.

